# Novel approach to quantitative spatial gene expression uncovers genetic stochasticity in the developing *Drosophila* eye

**DOI:** 10.1101/175711

**Authors:** Sammi Ali, Sarah A. Signor, Konstantin Kozlov, Sergey V. Nuzhdin

## Abstract

Robustness in development allows for the accumulation of neutral genetically based variation in expression, and here will be termed ‘genetic stochasticity‘. This largely neutral variation is potentially important for both evolution and complex disease phenotypes. However, it has generally only been investigated as variation exhibited in the response to large genetic perturbations. In addition, work on variation in gene expression has similarly generally been limited to being spatial, or quantitative, but because of technical restrictions not both. Here we bridge these gaps by investigating replicated quantitative spatial gene expression using rigorous statistical models, in different genotypes, sexes, and species (*Drosophila melanogaster* and *D. simulans*). Using this type of quantitative approach with developmental data allows for effective comparison among conditions, including health versus disease. We apply this approach to the morphogenetic furrow, a wave of differentiation that sweeps across the developing eye disc. Within the morphogenetic furrow, we focus on four conserved morphogens, *hairy, atonal, hedgehog*, and *Delta*. Hybridization chain reaction quantitatively measures spatial gene expression, co-staining for all four genes simultaneously and with minimal effort. We find considerable variation in the spatial expression pattern of these genes in the eye between species, genotypes, and sexes. We also find that there has been evolution of the regulatory relationship between these genes. Lastly, we show that the spatial interrelationships of these genes evolved between species in the morphogenetic furrow. This is essentially the first ‘population genetics of development’ as we are able to evaluate wild type differences in spatial and quantitative gene expression at the level of genotype, species and sex.

## Introduction

Natural genetic variation within populations has long been the purview of evolutionary and population geneticists, while developmental biologists focus on the effect of large mutations in otherwise isogenic backgrounds (Paaby and Gibson, 2016). This dearth of work on developmental variation in wildtype genetic backgrounds is in part because developmental approaches have long been restricted to data that is semi-quantitative (i.e. *in situ* hybridization, antibody staining). Indeed, gene expression studies are generally spatial or quantitative, but not both. In addition, the data is generally not interrogated from a quantitative perspective, including replication and rigorous statistical models. Without quantitative replication and statistical tests one cannot effectively compare developmental processes among conditions, including health versus disease. This is especially important given the potential for complex interactions between conditionally neutral differences in gene expression to result in disease phenotypes. Here we use hybridization chain reaction (HCR) to bridge this gap between developmental and quantitative or population genetics by quantitatively measuring spatial gene expression in multiple genotypes from two sexes of two species (*Drosophila melanogaster* and *D. simulans*) (Choi et al., 2014). Furthermore, we introduce rigorous replication allowing for statistical hypothesis testing. This is essentially the first ‘population genetics of development’ as we are able to evaluate wild type differences in spatial and quantitative gene expression at the level of genotype, species and sex. Multiplexing of four genes simultaneously also allows more rigorous analysis of gene co-expression, compared to other techniques that require inference across samples that have been individually stained. We use this enormous developmental dataset to focus on the well-known morphogens driving ommatidia specification in *Drosophila* (Atkins et al., 2013; W. Li et al., 1995; Raj et al., 2008; Shah et al., 2016; Tsachaki and Sprecher, 2011).

The *Drosophila* eye is formed from an imaginal disc, which is initially patterned by a wave of differentiation marked by a visible indentation of the tissue, termed the morphogenetic furrow (MF). The MF passes from the posterior to the anterior of the disc over a period of two days (90 minutes per adjacent row), giving each disc an element of both time and space in development (Fig 1) (Roignant and Treisman, 2009). A strength of the eye disc as a model is that within the MF all cells are arrested at G1, and there will be no additional variability introduced due to differences between cells in their stage of cell division (Baker and Yu, 2001; Escudero and M. Freeman, 2007; Firth and Baker, 2005; Firth et al., 2010). Furthermore, while there is evidence that transcription is bursty, the eye disc is composed of a repeated pattern of cells that will become ommatidia, akin to natural replication (Bothma et al., 2014; Fukaya et al., 2016; Tantale et al., 2016). By averaging across it, we reduce the impact of differences in transcriptional bursting across the eye disc.

**Figure 1.**
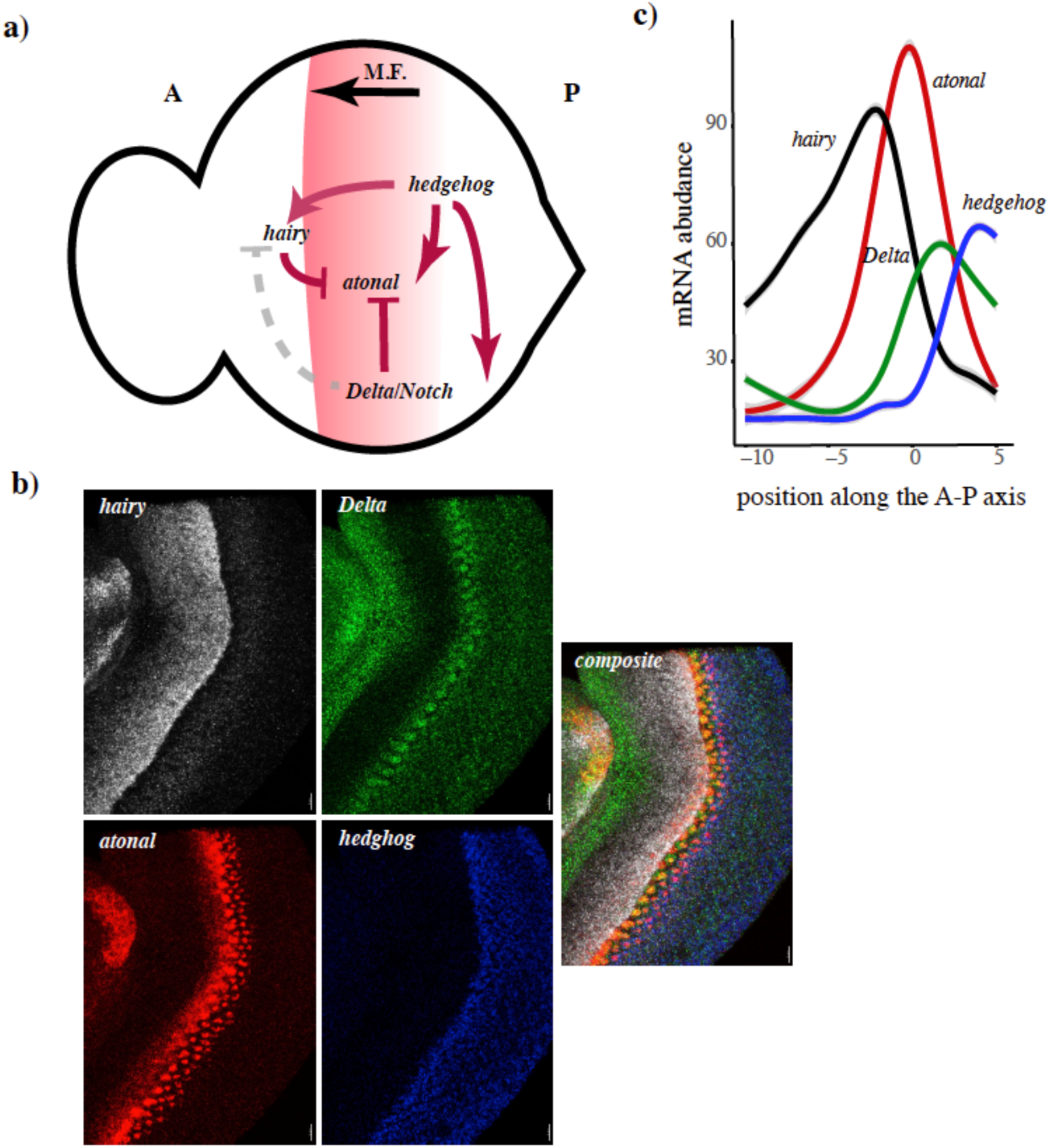
**a)** A summary of the eye patterning genes and pathway explored in this paper. The position of the MF is shown in red, and its direction of movement indicated below. Regulatory relationships are illustrated either as repression (bar) or activation (arrow). Regulatory relationships which are unclear are shown as gray dotted lines. **b)** Example images from the dataset, illustrating gene expression patterns of each gene. The composite image makes the additional point that we were able to analyze co-expression patterns of all four genes without needing to stain each gene in different samples and infer gene co-expression patterns. **c)** An illustration of the general expression pattern of each of the four genes in the study, along the anterior-posterior axis of the eye disc. The authors note that all curves shown in figures were plotted using smooth.spline in ggplot2 in R, and are meant only as visual representations of the data rather than formal reflection of the analysis.

The furrow is initiated by *hedgehog,* which both represses (short range) and activates (long range) *hairy* (Fig 1) (Felsenfeld and Kennison, 1995; Strutt and Mlodzik, 1997). *hairy* represses *atonal*, preventing precocious neural development anterior to the MF (though this role has been recently contested) (Bhattacharya and Baker, 2012; N. L. Brown et al., 1991; 1995). *hedgehog* activates the expression of *atonal*, driving the MF anteriorly (Fig 1) (Heberlein et al., 1993; Ma et al., 1993; Greenwood and Struhl, 1999). *atonal* is the proneural gene in *Drosophila*, establishing the competency to become photoreceptor cells (Jarman et al., 1994). The relationship between *Delta*/*Notch* and the other members of the pathway is more complex, but it is clear that in cells posterior to the furrow *Delta/Notch* repress *atonal* (Fig 1) (Firth and Baker, 2005; Gavish et al., 2016). There is also some evidence that *Delta*/*Notch* repress negative regulators of *atonal* at the furrow, such as *hairy* (Bhattacharya and Baker, 2009; N. L. Brown et al., 1995; A. B. A. M. Freeman, 2001). In addition, there is some evidence suggesting that *Notch/Delta* are involved in the early stages of *atonal* induction, and alternatively that *atonal* activates its own transcription (Baker and Yu, 1997; Dominguez and Hafen, 1997; Dominguez et al., 1998; Y. Li and Baker, 2001; Spencer et al., 1998; Sun et al., 1998). There are many other genes involved in the specification of the eye disc that will not be mentioned here, in favor of focusing on the genes we have assayed. We analyze the spatial quantitative expression of *hedgehog, hairy, atonal*, and *Delta* to understand the evolving regulatory logic of the gene network and changes in spatial dynamics between sexes and species.

We term differences in gene expression between species, genotypes, and sexes ‘genetic stochasticity’, as there are no phenotypic differences between these *Drosophila* eyes other than size and proportion of photoreceptor subtype. Size is included as a co-factor in the relevant models discussed below, and results only in a larger area being patterned for ommatidia rather than a difference in pattern. In addition, photoreceptor subtype is not determined or affected by the genes expressed during the initial patterning phase in the MF (Cook et al., 2003; Johnston and Desplan, 2014; Johnston et al., 2011; Wernet et al., 2006). We interpret genetic stochasticity in quantitative spatial patterns of gene expression in light of regulatory relationships among them. We approach a spatial and quantitative analysis of these gene expression patterns in three ways, first by explicitly creating a spatial gene expression profile and comparing between genotypes, sexes, and species. Second, we were interested in examining if the regulatory relationship between these genes had evolved between species or harbors variation within a species. Lastly, we investigated the possibility that the spatial relationship between these genes relative to the MF had evolved or harbors variation within populations.

## Methods

### Fly stocks

*D. simulans* were collected from the Zuma organic orchard in Zuma beach, CA in the spring of 2012 (Signor et al., 2017). They were inbred by 15 generations of full sib crosses. *D. melanogaster* were collected in Raleigh, North Carolina and inbred for 20 generations (Mackay et al., 2012).

### Staging and dissection of larvae

All flies were reared on a standard medium at 25° C with a 12-h light/12-h dark cycle. 120 hours after hatching, 3^rd^ instar larva were placed in phosphate buffered saline (PBS) and separated by sex. Their guts were carefully removed posteriorly and their body was inverted anteriorly to expose the brains, eye discs and mouth hooks. After fixation and labeling (described below), eye discs were isolated and mounted. The authors note that while there will be variation in the exact row of the eye that is being patterned between images, replicates were not conducted at the same time nor from the same cross. As such, with up to five replicates per line, any variation in the exact positioning of the furrow will serve to increase noise within the dataset rather than create false signal.

### Hybridization Chain Reaction (HCR)

HCR is unique in that it produces gene expression patterns that are both quantitative and spatial. The DNA probes were designed and synthesized by Molecular Instruments (Choi et al., 2014) (S1 Table). Four genes were multiplexed in each preparation as orthogonally-designed hairpins allowed the simultaneous amplification of their target sequences (Fig 1, S1 Fig). Each target mRNA was detected using five DNA probes to annotate the position and expression levels for each of the four assayed genes (*hairy, atonal, Delta and hedgehog*). Each probe contained two-initiator sequences (I1 and I2) that bound to a specific amplifier.

While other approaches such as FISH can be adapted to detect individual transcripts, HCR has a linear signal that is 20x brighter than FISH, it reduces non-specific background staining, and it can detect 88% of single RNA molecules in a cell with an appropriately low false discovery rate (Ma and Moses, 1995; Pan and Rubin, 1995). It is also highly repeatable, with different sets of probes targeted to the same gene showing correlations of .93-.99 (S. Fraser, pers. comm.).

The protocol for HCR was modified from (Choi et al., 2014) and is described briefly. The full protocol is available in S1 File. Inverted 3^rd^ instar larva were fixed in 4% paraformaldehyde diluted with PBS containing .2% Tween 20 (PBST). After fixation, larva were washed with PBST, then increasing concentrations of methanol (30%, 70% and 100%) at 25° C. Larva were stored in 100% methanol at −20° C. Methanol-fixed samples were thawed, washed with ethanol, re-permeabalized in 60% xylene, washed with ethanol, then methanol and rehydrated with PBST. Samples were permeabalized with proteinase K (4 mg/mL), fixed in 4% formaldehyde then washed with PBST at 25° C. Finally, at 45° C, samples were pre-hybridized for 2 hours before the addition of all the probes. The probe-hybridized larva were placed in wash buffer (Molecular Instruments) at 45° C to remove excess probes. Fluorescently labeled hairpins were snap-cooled then added to the samples at 25° C and placed in the dark to amplify the signal. Afterwards, samples were washed in 5X SSCT solution, isolated in PBST, then placed in Prolong Gold anti-fade mounting medium (Molecular Probes).

### Microscopy

Three dimensional images of mounted, HCR stained 3^rd^ instar larva eye discs were acquired on a Zeiss LSM 780 laser scanning microscope (Carl Zeiss MicroImaging, Inc., Thornwood, NY, USA) with Objective Plan-Apochromat 63x/1.40 Oil. The gain was adjusted to avoid pixel saturation.

### Extraction of gene expression profiles

The first steps in the image analysis is bringing each image to the same orientation and segmenting it. Image segmentation produces a mask in which pixels are assigned to objects or background. Here the objects are one or several mRNA molecules. Then the cellular structure of the imaginal disc is approximated using a hexagonal array. Though the real underlying cell structure of the imaginal disc is technically able to be recognized, this was unsatisfactory in our data due to imaging noise. Thus, at the second step using the R package hexbin we constructed a partition of the imaginal disc area into elements that represent pseudo-cells and have a biologically-relevant hexagonal shape (Brennan et al., 1998). The number of pseudo-cells was selected by visual inspection of the combined image in which the hexagonal structure was overlaid onto the *atonal* channel to verify fit. We are primarily interested in expression profiles around the MF, providing us a convenient landmark to align images from different preparations, thereby assigning coordinates to the pseudo-cells. However, deformations of the eye disc during growth and preparation sometimes distorts the MF. We used splines to correct for any bending or deformation of the MF. Next, using the histograms of cumulative pixel intensities of objects in expression domains and non-expressing areas we estimated the typical intensity of a transcript and typical background signal, respectively. Consequently, the cumulative intensities greater than the background are divided by the intensity attributed to single mRNA molecule to yield counts of mRNA molecules. This normalizes the expression profiles and corrects for differences in microscope gain between images. Finally, the gene expression profiles are estimated for every pseudo-cell.

### Morphological reconstruction and contrast mapping segmentation

To detect gene transcripts within the image stacks we applied a version of the MrComas method that was modified for processing 3D images (Kozlov et al., 2017). This approach first enhances contrast within the image and reduces noise. The images were enlarged by a factor of four with the nearest-neighbor algorithm. They were processed by morphological reconstruction using both opening and closing, where closing (opening) is dilation (erosion) that removes extraneous dark (bright) spots and connects bright (dark) objects (Vincent, 1993). The contrast mapping operator assigns each pixel the maximum value between the pixel-by-pixel difference of the reconstructed images and their pixel-by-pixel product and produces the rough mask for each channel. An image, *I*, is mapping from a finite rectangular subset *L* onto the discrete plane *Z*^2^ into a discrete set 0, 1,…, N − 1 of gray levels. Let the dilation *δB* and erosion *ϵB* by structural element *B* be defined as:

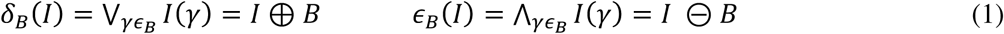

Where V and ∧ denote infimum and supremum respectively. Then formulae:

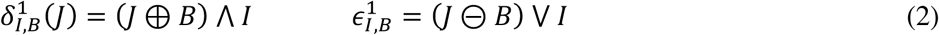

denote geodesic dilation 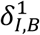 and erosion 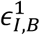. Binary reconstruction extracts those connected components of the mask image which are marked on the marker image, and in grayscale it extracts the peaks of the masked image marked by the marker image. Using the dilated masks image *I* as the marker *J*: *J* = *δ*_*B*_(*I*) defines closing by reconstruction:

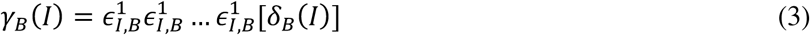

Opening by reconstructions uses eroded mask *I* as a marker *J*:

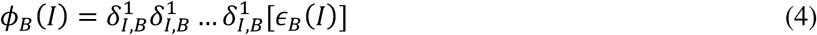

Then the difference between closing and opening by reconstruction has the meaning of the gradient:

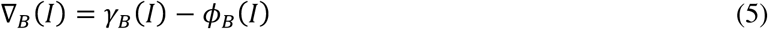

To create strong discontinuities at object edges and flatten signal with the objects the contrast mapping operator takes a maximum between the difference and the pixel-by-pixel produce of the reconstructed images and produces a rough mask for each channel:

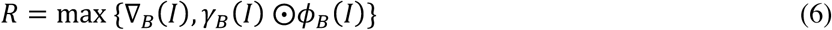

Subsequently, this mask was subjected to distance transform, which substituted each pixel value with the number of pixels between it and the closest background pixel. This operation creates ‘peaks’ and ‘valleys’ of intensity inside foreground objects. To split erroneously merged objects watershed transform was applied, which treats the whole image as a surface and intensity of each pixel as its height and determines the watershed lines along the tops of ridges separating the catchment basins (Meyer, 1994). The quality of segmentation is assessed visually by inspection of the object boarders overlaid with the original image. Finally, each mask is returned to its original size and quantitative measures are made of shape and intensity characteristics such as the number of pixels, as well as their mean and standard deviation in the detected object. MrComas is free and open source software available at http://sourceforge.net/p/prostack/wiki/mrcomas.

### Approximating the MF

We defined the position of the MF as the middle of overlap between *hairy* and *atonal* expression. The shape of the MF was approximated with a spline using function smooth.spline in R. The degrees of freedom and other parameters were chosen to make the approximation coincide visually with a MF image.

### Inferring counts of transcript number

Segmentation of the image provided a table of coordinates and the shape and intensity characteristics of detected transcripts. Here, we applied filtering steps to remove false positives and determine the count of mRNA transcripts. First, we assumed the object we detected as least intense but most frequent corresponds to a single mRNA molecule. Then, we inferred background intensity for objects outside of well-annotated domains of expression of the four genes. Assuming that the majority of true objects contain a single molecule, we compare the distribution of cumulative intensities of particles in expression domains and areas of known non-expression to obtain the typical intensity of a true single molecule and a false positive, respectively. All detected signals that were lower than the typical intensity of a false positive were removed from the dataset. The number of removed objects is typically less then 10%. All other cumulative pixel intensities were divided by the typical intensity of a true single molecule as normalization coefficient to yield an estimate of the number of mRNA transcripts.

### Image registration

We applied an affine coordinate transformation to each eye disc to make the corresponding maxima and the width of expression patterns of four genes in different eyes coincide as closely as possible. To do so, we shifted the coordinate system of each eye to its center and also scaled them in the A-P direction. The center of the pattern in A-P direction is the MF.

We mapped the expression patterns to a unified hexagonal structure in order to make comparisons between pseudo-cells from individual imaginal discs. The unified cell structure was constructed using the R package hexbin. Each cell in the unified grid represents an ‘average’ cell from individual eyes. The size of a hexagon in the unified grid is greater or equal than the cell size in the individual eye. Thus, the number of molecules in each unified cell in the mapped pattern equals the mean over the cells from native pattern that are covered by this unified cell. After such coordinate transformation, the MF region is defined as 20 cells on either side of the MF, to focus the analysis on the area of interest (the MF).

### Filtering and quality controls for each eye disc

Some eye imaginal discs were damaged or deformed in the process of dissection or mounting, resulting in regions of erroneous gene expression, such as disruptions to the MF. The expression profile of each disc was examined by eye and these regions were individually trimmed out of the final dataset. At the edges of each eye disc the pattern of the MF was also degraded, so each eye disc was trimmed dorso-ventrally prior to analysis. Outliers were excluded from the dataset, determined as a single member of the five replicates with more than a 3x difference in expression values. This resulted in a final dataset of 55 eye discs.

### Analysis of individual spatial gene expression patterns

We were primarily interested in variation in gene expression profiles across the eye disc, that is using differences in expression averaged across rows along the *x*-axis. While the *y*-axis is of interest, variation in the shape, size, degree of deformation, and occasional damage to the disc made this analysis intractable. We fit curves to each gene expression profile using the mgcv package in R, using a generalized additive model with integrated smoothness estimation. Smoothing terms are represented using penalized regression splines. predict.gam was used to fit the curves to the original range of values and down sample the curves to eight points. MANOVAs were performed using the “Pillai” test for species × genotype × sex.

### Modeling framework to understand variation and evolution of the eye patterning gene network

We wanted to understand if cryptic variation existed within the regulatory logic of *hairy, atonal, Delta*, and *hedgehog*, or if there had been cryptic evolution between species. To understand the regulatory logic between genes we focused on biologically relevant relationships, such as the regulation of *hairy* by *Delta* and *hedgehog*, but excluded such relationships as *hairy* and *atonal*. This was due to the low overlap between *hairy* and *atonal* expression domains, where including cells where only one or another was expressed would artificially create a relationship between expression levels. Both *hairy* and *atonal* are downstream, directly or indirectly, of *Delta* and *hedgehog* thus it was these relationships that were modeled. We limited the analysis to cells where all genes included were expressed in at least ten molecules.

In the previous analysis, we investigated variation in the cryptic spatial quantitative expression pattern of genes in the MF. Here, we will investigate the possibility that genes in the MF have evolved, or harbor variation, for how they affect each other in particular cells. For example, is high *atonal* expression associated with high expression of *hedgehog*, given that *hedgehog* activates *atonal*? We used the following equations to determine the relationship between the expression of these genes:

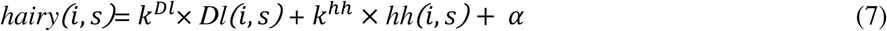

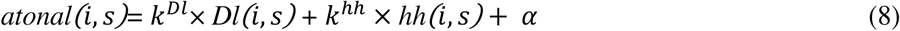

The coefficient *k* and constant *α* were fit using standard methods for multiple regression. Here *hairy(i, s)* and *atonal(i, s)* are the measured expression level of each gene in cell *i* in individual *s*. *Dl(i, s)* and *hh(i, s)* are vectors containing the corresponding expression levels of *hairy* and *atonal*‘s regulators *Delta* and *hedgehog*. To determine if the regulatory logic is the same between genotypes and species we can then use the regression coefficients from these models in a MANOVA. We note that we cannot exclude the possibility that other unmeasured genes are responsible for producing this variation.

### Model for understanding overall variation and evolution of MF structure

Lastly, we wanted to understand if there is variation in the relationship between the MF and gene expression, or if variation existed for the size of the MF overall. For example, the MF was called as the position of overlap between *atonal* and *hairy* expression, but it is unclear how the overall gene expression pattern of these genes relates to their overlap. For example, is the position of maximum expression of each always the same relative to the MF? Two processes occurred to make the MF comparable between samples, they were shifted to occupy the same position depending upon the position of overlap of *hairy* and *atonal*, and they were scaled to occupy the same total area. The amount required to scale will depend both on the size of the original disc and the width of the MF relative to the disc. To account for differences in size we include the number of rows in the original disc prior to any transformations as a cofactor and perform ANOVA in R.

## Results

### Individual spatial gene expression patterns

First, to characterize the spatiotemporal dynamics of transcriptional activity along the anterior-posterior axis, we took the spatial average of signal across the dorsal-ventral axis and compared between genotypes, sexes, and species (Fig 2A). The authors note that smoothed curves in the figures were created using smooth.spline in ggplot2, which is not the same method for curve fitting as in the analysis. As such they are meaningful reflections of the patterns in the data but not depictions of actual analyses. We found abundant spatial quantitative variation in expression profiles (Fig 2-3). The expression profile of *hairy* around the MF harbors variation between genotypes and there is an interaction between genotype and sex (Table 1, *p* = 2 × 10^−3^, *p* = 0.02). There has also been evolution between species for *hairy* (Table 1, *p* = 3 × 10^−4^). While *atonal* has not evolved between species, there is variation in expression profile between genotypes, sexes, and there is an interaction between genotype and sex (Fig 3A-C, Table 1, *p* = 4 × 10^−4^, *p* = .02, *p* = .02). Surprisingly, given the conservation of *Delta* in general, *Delta* harbors variation in spatial quantitative expression behind the MF between genotypes and sexes (Table 1, *p* = 2 × 10^−3^, *p* = 7 × 10^−4^) and there are significant interactions between genotype and sex (Fig 2B-C, Table 1, *p* = 2 × 10^−3^). There has also been evolution of *Delta* between species, and evolution of the interaction between species and sex (Table 1, *p* = .03, *p* = 3 × 10^−4^). *hedgehog* is not different between species but is significantly different between genotypes, sexes, and there is an interaction between the two (Table 1, *p* = 5 × 10^−4^, *p* = .05, *p* = 1 × 10^−3^). There is also a significant interaction between species and sex (Table 1, *p* = .01).

**Figure 2.**
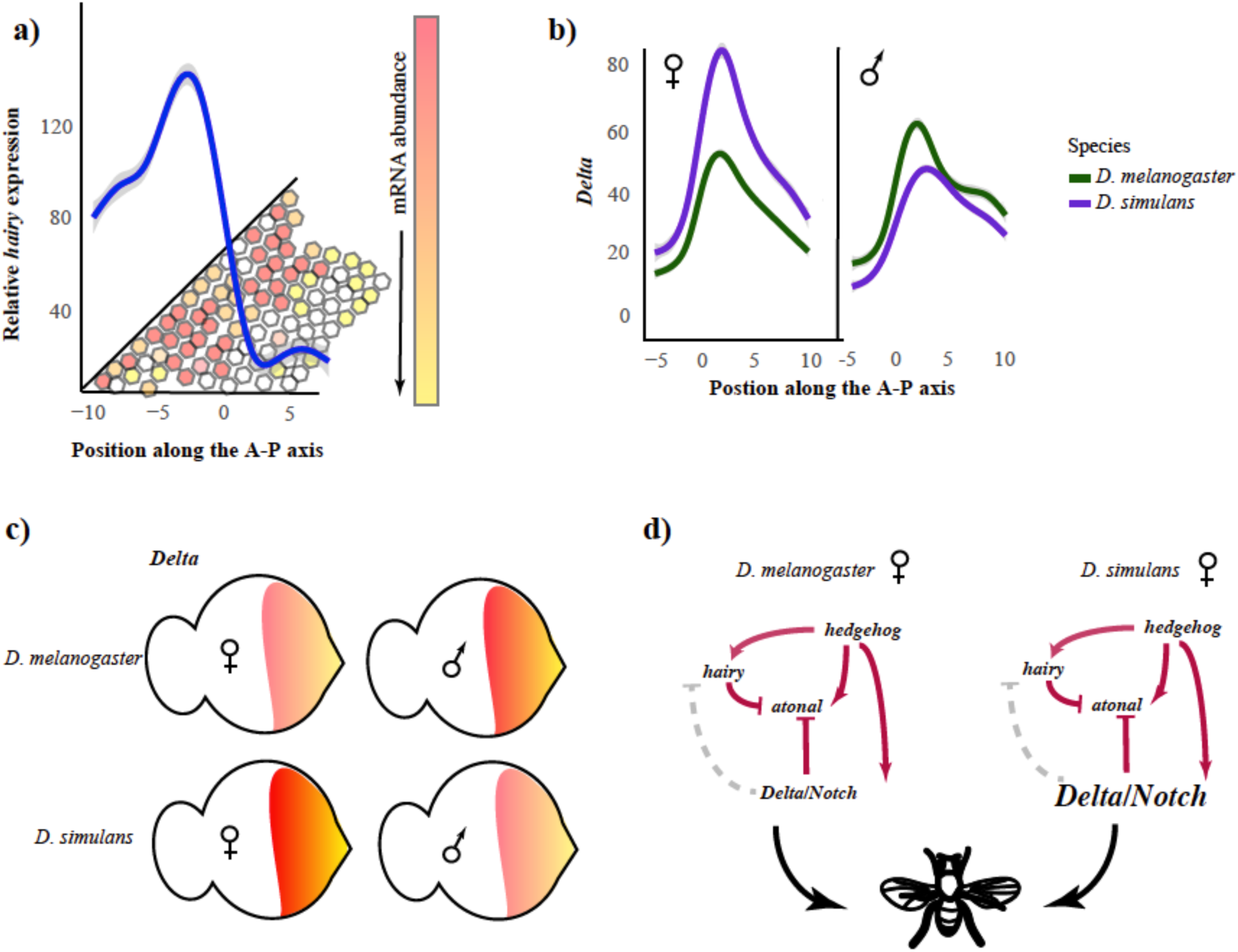
**a)** This is an example of a curve being fitted to the gene expression profiles, though note that the curve corresponds to the average in a given row (*x*-axis). The hexagons are intended to represent cells with varying amounts of *hairy* expression, from the highest (red) to the lowest (white). **b)** An illustration of variation in *Delta* expression between species and sexes. Curves shown are fitted to all genotypes within a sex and species with confidence intervals indicated in gray. **c)** An illustration using the imaginal disc of how *Delta* expression varied between species and sexes, with lower expression in *D. melanogaster* females and *D. simulans* males **d)** Evolution of *Delta* illustrated within the context of the gene network, illustrating how changes in *Delta* expression are not perturbing the gene network and result in phenotypically normal *Drosophila*.

**Figure 3.**
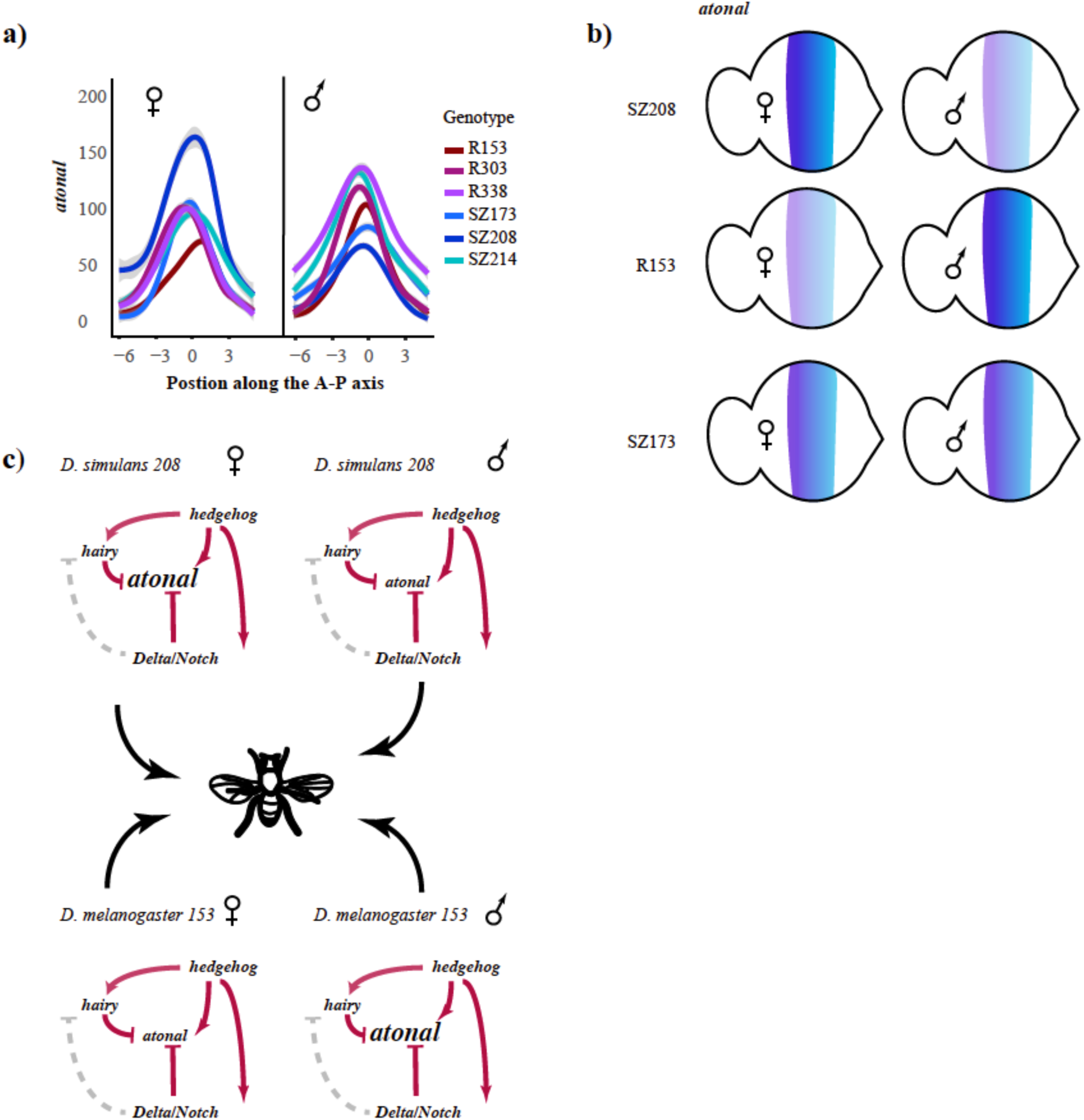
**a)** An illustration of variation in *atonal* expression between genotypes and sexes. Curves shown are fitted to each genotype and sex with confidence intervals indicated in gray. **b)** An illustration using the imaginal disc of how *atonal* expression varied between genotypes, with lower expression in females of *D. melanogaster* R153 and males of *D. simulans* Sz208. *D. simulans* Sz173 has lower expression than females of Sz208 but it is not sexually dimorphic. **c)** Evolution of *atonal* illustrated within the context of the gene network, illustrating how changes in *atonal* expression are not perturbing the gene network and result in phenotypically normal *Drosophila*.

**Table 1.** The results of the full model for each gone.

| Effect | numDF | denDF | F-value | p-value |
| --- | --- | --- | --- | --- |
| Species | 8 | 35 | 5.1 | **$3 \times 10^{-4}$** |
| Genotype | 32 | 152 | 2.07 | **$2 \times 10^{-3}$** |
| Sex | 8 | 35 | 1.08 | 0.32 |
| Species x Sex | 8 | 35 | 1.94 | 0.08 |
| Genotype x Sex | 32 | 152 | 1.73 | **0.02** |

*atonal*
| Effect | numDF | denDF | F-value | p-value |
| --- | --- | --- | --- | --- |
| Species | 8 | 35 | 1.53 | 0.18 |
| Genotype | 32 | 152 | 2.31 | **$4 \times 10^{-4}$** |
| Sex | 8 | 35 | 2.6 | 0.02 |
| Species x Sex | 8 | 35 | 1.91 | 0.09 |
| Genotype x Sex | 32 | 152 | 1.73 | **0.02** |

*Delta*
| Effect | numDF | denDF | F-value | p-value |
| --- | --- | --- | --- | --- |
| Species | 8 | 35 | 2.43 | **0.03** |
| Genotype | 32 | 152 | 2.07 | **$2 \times 10^{-3}$** |
| Sex | 8 | 35 | 4.55 | **$7 \times 10^{-4}$** |
| Species x Sex | 8 | 35 | 5.03 | **$3 \times 10^{-4}$** |
| Genotype x Sex | 32 | 152 | 2.03 | **$2 \times 10^{-3}$** |

*hedgehog*
| Effect | numDF | denDF | F-value | p-value |
| --- | --- | --- | --- | --- |
| Species | 8 | 35 | 0.87 | 0.55 |
| Genotype | 32 | 152 | 2.28 | **$5 \times 10^{-4}$** |
| Sex | 8 | 35 | 2.23 | 0.05 |
| Species x Sex | 8 | 35 | 2.88 | 0.01 |
| Genotype x Sex | 32 | 152 | 2.17 | **$1 \times 10^{-3}$** |

Significant *p*-values are indicated in bold, with gray shading.

Thus, *hairy* and *Delta* have evolved different spatial quantitative expression patterns between species, while *hairy, atonal, Delta*, and *hedgehog* harbor cryptic variation within species and sexes. Given that there are regulatory relationships between these genes, it is interesting to see that they do not all harbor variation for the same factors. This could potentially be due to the influence of other unmeasured regulatory factors, or to variation in the relationship between these genes and other components in the gene regulatory network. However, whatever the source of ‘buffering’ of the network, be it the effect of other genes or threshold effects on development, the fact that this information is not retained within the steps of the pathway supports our supposition that this variation does not ultimately have a phenotypic effect.

### Variation and evolution of the eye patterning gene network

There has been evolution in the regulatory logic of *hairy* and its upstream regulators *Delta* and *hedgehog* between species (Fig 4A-C, Table 2, *p* = .03). There is also variation between sexes in the regulatory logic of *hairy* and its upstream regulators *Delta* and *hedgehog* (Table 2, *p* = .03). There has been significant evolution of the regulatory logic of *atonal*, in a significant interaction between species and sex (*p* = 1 × 10^−3^). Furthermore, while there was no significant effect of genotype for *hairy*, there is for *atonal*, indicating that there is variation segregating in the population affecting the relationship between *atonal, hedgehog*, and *Delta* (*p* = 1.6 × 10^−5^). There is also a significant interaction between genotype and sex (*p* = 1 × 10^−3^). Thus, the relationship between *hairy* and *atonal* and their regulators has evolved between species and sexes in *hairy*, and between genotypes and sex in *atonal*. We illustrate this difference between species in Figure 3, where a different relationship between *hairy* and *hedgehog* is visible between *D. melanogaster* and *D. simulans*. In brief, the frequency of cells with a given log transformed level of expression are plotted against one another for *hairy* and *hedgehog*. *hairy* is primarily expressed anterior to the MF and *hedgehog* posterior, and they have a different regulatory relationship in each region with *hedgehog* activating *hairy* long range (anterior) and repressing it short range (posterior). This is reflected in the frequency of cells expressing both genes for *D. melanogaster*, where anterior to the MF there is a high frequency of *hairy* expressing cells and a low frequency of co-occurring high *hedgehog* expression. Posterior to the MF the opposite is true, with high expression of *hedgehog* lacking concordance with any expression of *hairy*. In *D. simulans*, posterior to the MF, this relationship is the same as in *D. melanogaster*. However, in anterior to the MF this is not the case. Expression of *hairy* and *hedgehog* both increase as the other increases, with widespread co-occurrence.

**Figure 4.**
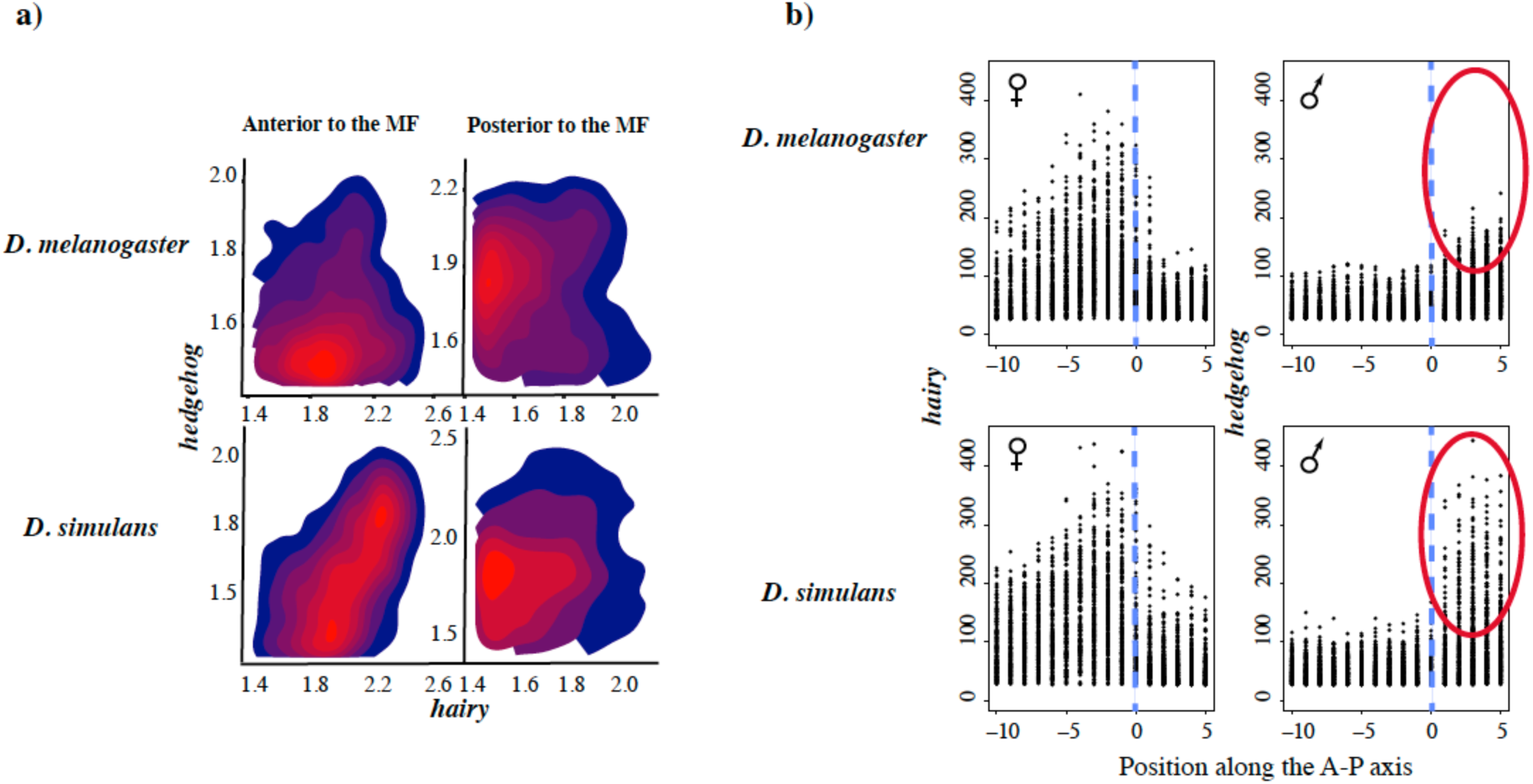
**a)** An example of variation in regulatory logic between *D. simulans* and *D. melanogaster* for *hairy* and *hedgehog*. The heat map illustrates the density of points, and thus reflects the frequency of a given co-expression profile between *hairy* and *hedgehog*. Gene expression values were log-transformed to better illustrate lower values and split between anterior to the MF and posterior to the MF. The split between the two regions was to investigate the possibility that *hedgehog* had a different regulatory relationship with *hairy* depending upon its relationship to the MF, given that *hedgehog* is thought to activate *hairy* long range and repress *hairy* short range. **b)** An illustration of the change in quantitative spatial expression of *hairy* and *hedgehog* between species and sexes, with the position of the center of the MF marked with a dotted blue line. The red circle emphasizes a large change in maximum *hedgehog* expression in males of the two species.

**Table 2:** The results of the full model for regulatory relationship between *hairy*, *Delta*, and *hedgehog*, and *atonal*, *Delta*, and *hedgehog*.

| Effect | numDF | denDF | F-value | p-value |
| --- | --- | --- | --- | --- |
| Species | 2 | 41 | 4.03 | **0.25** |
| Genotype | 8 | 84 | 1.2 | 0.31 |
| Sex | 2 | 41 | 3.95 | **0.027** |
| Species x Sex | 2 | 41 | 0.78 | 0.46 |
| Genotype x Sex | 8 | 84 | 1.11 | 0.37 |

***atonal***
| Effect | numDF | denDF | F-value | p-value |
| --- | --- | --- | --- | --- |
| Species | 2 | 41 | 2.64 | 0.08 |
| Genotype | 8 | 84 | 5.43 | **$1.6 \times 10^{-5}$** |
| Sex | 2 | 41 | 0.67 | 0.52 |
| Species x Sex | 2 | 41 | 8.09 | **$1 \times 10^{-3}$** |
| Genotype x Sex | 8 | 84 | 2.41 | **0.02** |

Significant *p*-values are indicated in bold, with gray shading.

### Variation and evolution of MF structure

The amount that the eye discs were shifted is not significant for genotype, sex, or species, suggesting that the relationship of maximum gene expression with the MF does not vary. However, the amount that they were scaled is, after accounting for original differences in size, between species (*p* = 1.38 × 10^−6^). This suggests that the total relative width of the MF varies between species, but not between genotypes or sexes. This is also suggestive of evolving interrelationships among genes that could result in broader or narrower areas in which they enhance or suppress expression of one another.

### Discussion

Our results summarize a complicated pattern of variation sorting in the gene network involved in patterning the MF. For example, the overall shape of the expression of *hedgehog* across the eye disc is different between genotypes, sexes, and there is an interaction between species and sex and genotype and sex. *hedgehog* upregulates *hairy*, but *hairy* has differences in expression between species (which *hedgehog* does not), genotypes, and there is an interaction between genotype and sex. Thus, the differences seen in upstream regulators, such as *hedgehog*, are not recapitulated in their downstream targets. In another example, *Delta/Notch* is expected to repress *atonal*, but while *Delta/Notch* is significant for all categories tested *atonal* is only significant for genotype, sex, and their interaction. It is possible that this variation is being mitigated or dampened by other regulatory factors not assayed here, or that certain aspects of genetic background are more or less sensitive to variation. For example, that fixed variation between species dampens variation at *Delta/Notch* but sorting variation remains sensitive between genotypes, which propagates to *atonal*.

It may be that all of this variation is within levels tolerated by the network, as it has been shown that gene networks can have thresholds of variation, below which differences in expression are effectively neutral. These thresholds can also be two sided, creating a sigmoidal curve the center of which is neutral phenotypic space (Felix and Barkoulas, 2015). Many studies have shown a relative insensitivity to variation in gene dosage, for example in *Drosophila* early embryos the *bicoid* gradient results in normal development at one to four dosages of the gene, but markedly abnormal development at six or more (Liu et al., 2013; Lucas et al., 2013; Namba et al., 1997). It is also possible that the ‘genetic stochasticity’ documented in these genes is in fact deleterious, and is being compensated for elsewhere in the network. While most deleterious mutations are purged by selection, they may rise in frequency due to genetic drift or hitchhiking, among other possible causes (Burch and Chao, 1999; Chun and Fay, 2011; Estes and Lynch, 2003; McKenzie and Clarke, 1988). This type of compensatory mutation has been documented in microbial and animal systems (K. M. Brown et al., 2010; Burch and Chao, 1999; Charusanti et al., 2010; Estes and Lynch, 2003; Estes et al., 2011; Maisnier-Patin and Andersson, 2004; McKenzie, 1993; McKenzie et al., 1982; Moore et al., 2000; Stoebel et al., 2009; Szamecz et al., 2014). Recently cell cycle heterogeneity has been implicated in the appearance of widespread noise in development, however this is not responsible for the genetic stochasticity observed here as all MF cells are arrested at G1 (Keren et al., 2015; Kumar, 2013).

There have been other semi-quantitative approaches to studying spatial gene expression patterns. In another study on *orthodenticle*, the authors found that the spatial and temporal pattern of gene expression was conserved but the amount of gene product was not, though this work was not strictly quantitative given that measurements were from *in situ* hybridization and reporter constructs and there was no rigorous statistical testing (Goering et al., 2009). This is in contrast to our results which showed significant differences in the spatial relationship between gene expression patterns between species. Other semi-quantitative works on the *Drosophila* embryo using *in situ* hybridization found that the regulatory relationship between genes in the anterior-posterior blastoderm patterning network were conserved, despite differences between species in their spatio-temporal pattern (Fowlkes et al., 2011; Wunderlich et al., 2012). Here we find that the regulatory relationship between *atonal* and *hairy*, and their regulators *hedgehog* and *Delta*, has evolved between species, sexes, and genotypes.

One of the important messages from this work is that rigorous statistical testing can uncover molecular variation in spatial and quantitative developmental gene expression. Using the type of replication applied in quantitative genetics with developmental data we were able to apply rigorous statistical models to micro-evolutionary variation in development. Despite this variation, observed with repeatable observations of developmental patterns among natural genotypes, the phenotypes of all flies are normal. This points to a potential abundance of hidden noise in spatial and quantitative gene expression. The evolutionary approach to development generally targets large changes that have occurred over broad phylogenetic distances (Ito et al., 2013; Jeong et al., 2008; Kopp et al., 2000; Reed et al., 2011; Rosenblum et al., 2010; Signor et al., 2016; Yassin et al., n.d.). Accordingly, the presence of abundant underlying variation is perhaps not a huge surprise. But it does have large implications, as it demands that we modify developmental models in such a way that such abundant genetic variation is buffered from perturbing the final phenotype. In the future, application of this type of replicated, quantitative, spatially resolved data will have unique insights into the penetrance of disease phenotypes and the origin of developmental defects.

## Data Availability Statement

All data associated with the manuscript can be found at:

## Author contributions

S. A. performed the staining and imaging of the eye discs, K. K. processed the imaging data, S.S. conceived of the experiment, analyzed the processed data and wrote the paper, S. N. conceived of the experiment and coordinated the research.

## Acknowledgements

The authors thank S. Restrepo, M. Samsonova, A. Kopp, P. Marjoram, J. Butler, G. Mackerel, R. Hudson, E. Williams, A. Lana, S. G. Frusina, H. Cagni and R. Hess for help with experimental procedures and manuscript preparation. This work was supported by grants U01GM103804 and RO1GM102227 to S.V.

## Supporting Information

**S1 Fig. Typical anterior-to-posterior expression patterns in the MF.** *hairy* (white), *atonal* (red), *Delta* (green), and *hedgehog* (blue), 3rd instar larva eye disc labeled using HCR. Please note that these images contain portions of the antennal disc, which were excluded from any analysis. Scale bars are 30 µm.

**S1 File Protocol: Hybridization chain reaction *Drosophila* imaginal discs**

**S1 Table The sequence of each of the probes used for hybridization chain reaction.**

